# Combining eye tracking with EEG: Effects of filter settings on EEG for trials containing task relevant eye-movements

**DOI:** 10.1101/2020.04.22.054882

**Authors:** Louisa Kulke, Vincent Kulke

## Abstract

Co-registration of electroencephalography (EEG) and eye movements is becoming increasingly popular, as technology advances. This new method has several advantages, including the possibility of testing non-verbal populations and infants. However, eye movements can create artefacts in EEG data. Previous methods to remove eye-movement artefacts, have used high-pass filters before data processing. However, the role of filter settings for eye-artefact exclusion has not directly been investigated. The current study examined the effect of filter settings on EEG recorded in a dataset containing task-relevant eye movements. Part 1 models the effects of filters on eye-movement artifacts and part 2 demonstrates this effect on an EEG dataset containing task-relevant eye-movements. It shows that high-pass filters can lead to significant distortions and create artificial responses that are unrelated to the target. In conclusion, high-pass filter settings of 0.1 or lower can be recommended for EEG studies involving task-relevant eye movements.

**Highlights:**

- Co-registration of EEG and eye-tracking is gaining popularity
- However, eye movements can create artifacts in the EEG signal
- The current paper models the effect of high pass filters on eye-movement artifacts
- High pass filters can induce large distortions in EEG data containing regular eye-movements
- The distortion is affected by fixation duration and filter frequency

## Introduction

Combined eye tracking and EEG is becoming more popular due to recent advances of technology (e.g. Dimigen, 2014; Kamienkowski, Ison, Quiroga, & Sigman, 2012; Kulke, Atkinson, & Braddick, 2015; Kulke & Wattam-Bell, 2013; Langer et al., 2017; Meyberg, Werkle-Bergner, Sommer, & Dimigen, 2015). The co-registration makes it possible to test non-verbal populations, like infants (Kulke, Atkinson, & Braddick, 2016b) and has been used in different areas of research (see e.g. Fischer, Graupner, Velichkovsky, & Pannasch, 2013; Huber-Huber, Ditye, Fernández, & Ansorge, 2016; Kulke, 2019; Kulke, Atkinson, & Braddick, 2016a; Kulke et al., 2016b; Nikolaev, Jurica, Nakatani, Plomp, & van Leeuwen, 2013; Nikolaev, Nakatani, Plomp, Jurica, & van Leeuwen, 2011).

Tasks combining these methods often involve eye movements as a fixed constituent of all trials. As eyes have an innate polarity, being positive at the side of the cornea and negative at the side of the retina (Jervis, Ifeachor, & Allen, 1988), eye-movements cause a change in electrical potentials measured at the scalp surface. It is therefore generally recommended to exclude any trials from EEG analyses that contain eye-movements as artefacts (see e.g. Jervis et al., 1988; Luck, 2005). However, when studying overt attention shifts, each trial is bound to contain an eye-movement, making it impossible to exclude these trials.

One way to avoid confounds of EEG with eye-movement artefacts, is to identify eye-movement-related components using independent component analysis (ICA). Although this method has been shown to successfully remove eye artefacts (Makeig, Bell, Jung, & Sejnowski, 1996; Mannan, Kim, Jeong, & Kamran, 2016; Plöchl, Ossandón, & König, 2012), neural responses that coincide with eye-movements might also be filtered out by the algorithm (Nikolaev, Meghanathan, & van Leeuwen, 2016; Plöchl et al., 2012). A recent paper by Dimigen (2020) suggests that ICA-based removal of eye-movement artefacts can be successfully used to remove artefacts but unfortunately also introduces distortions due to overcorrection of the data. Furthermore, the signal during overt attention shifts contains components induced by the change in visual scene due to eye-movements, which can lead to visually evoked potentials in response to the change rather than the initial stimulus.

Alternatively, stimulus onset related eye-movement artefacts can be avoided by extracting only short intervals (180 ms), so that most eye-movements occur after the extracted time window (Kulke, 2019; Kulke et al., 2015, 2016a, 2016b; Kulke, Atkinson, & Braddick, 2020). However, guides to EEG methods recommend digital filters to be applied to the EEG data before these time windows are extracted to avoid edge artefacts (e.g. Luck, 2005; Tanner, Morgan Short, & Luck, 2015), meaning that the data still includes eye-movement-related components when the filter is applied. Previous research suggests that high pass filters of 0.3 Hz and above can lead to artificial responses of opposite polarity preceding an actual response (Tanner et al., 2015) in data without task-relevant eye-movements. However, task-relevant eye-movements induce particularly large responses, potentially leading to even larger artefacts than are present in data without task-relevant eye-movements. This study aimed to investigate the effects of high-pass filters on EEG data containing regular eye-movements.

### Part 1: Modelling

The aim of this first part was to model the effect of eye-movements on EEG data. The eye movement was considered as a rectangular signal with a width of 0.7 s, the fixation duration used in several gaze-contingent studies (Kulke, 2019; Kulke et al., 2016a, 2016b), to model how it can lead to potential filter artefacts. When a high-pass filter (e.g. Butterworth Filter 3^rd^ order, filter frequency of 0.1 Hz) is applied to such a signal, the lower frequencies are blocked or weakened. In both the filtered and unfiltered signal, the frequency spectrum remains the same except for low frequencies that are affected by the high-pass filter. The signal in the frequency domain can be transformed back into the time domain using an inverse Fast Fourier Transformation (FFT). In the time domain, the lack of low-frequency oscillations due to the high-pass filter affects slopes in the vicinity of the rectangular pulse. The higher the filter frequency, the more low-frequency oscillations are removed from the signal. As a result, the filter error increases and attenuates the signal, which is expressed in a higher slope before and after the rectangular pulse (Figure 1).

**Figure 1.**
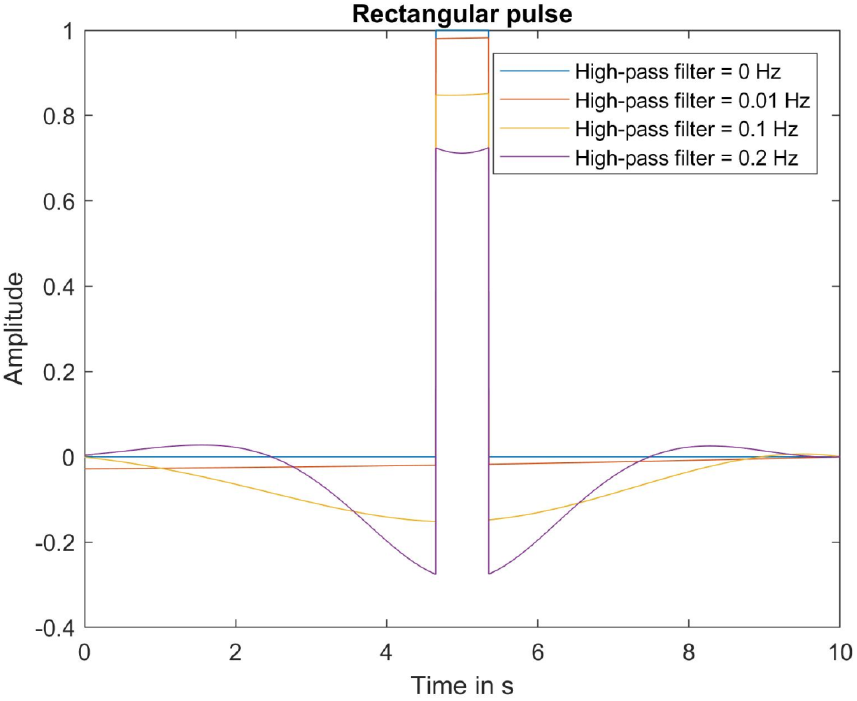
Time signal of a rectangular pulse with the filter error after inverse FFT

To quantify this error, the mean slope from 0.2 s before the rectangular pulse (approximate time period during which ERPs can be investigated) up to the beginning of the pulse can be computed, as a function of different parameters. The error strongly depends on, firstly, the width of the signal in the time domain. The frequency distribution of a rectangular impulse can be separated into harmonic blocks, with the first harmonic block having the highest amplitude, hence contributing most to the signal. The wider the signal is in the time domain, the smaller the harmonic blocks become in the frequency domain. If these blocks are smaller, in particular the first harmonic block, the filter will more strongly affect the signal, as a higher proportion of the first harmonic block will be removed or weakened through the filter. For example, if the filter frequency is 0.1 Hz and the main amplitude of the signal lies under 1 Hz, 10% of the main signal (not counting the rather small higher harmonics) is blocked or weakened. However, if the main amplitude of the signal lies between 0 and 10 Hz, only 1% of the signal is blocked or weakened; therefore, the error is considerably smaller. Secondly, the slope depends on the filter frequency. The higher the high-pass filter frequency, the greater the proportion of the signal that is cut off and hence the greater the artifactual slope. Figure 2 shows the influence of filter frequency and pulse width (which is directly related to the width of the signal in the frequency domain) on the slope (filter error). The pulse amplitude (normalised below) has a negligible impact on the slope. The model shows that the optimal filter frequency depends on the signal. In general, a lower filter frequency and a smaller pulse width result in a smaller absolute error. However, at a very high filter frequency, the average absolute slope is slightly reduced with increasing pulse width, contradicting the overall pattern. Note, however, that this does not imply that the artefact is reduced. In contrast, for extremely high values, such a high percentage of the spectrum is blocked or at least weakened that the resulting signal is highly deformed and the original signal is no longer recognizable.

**Figure 2.**
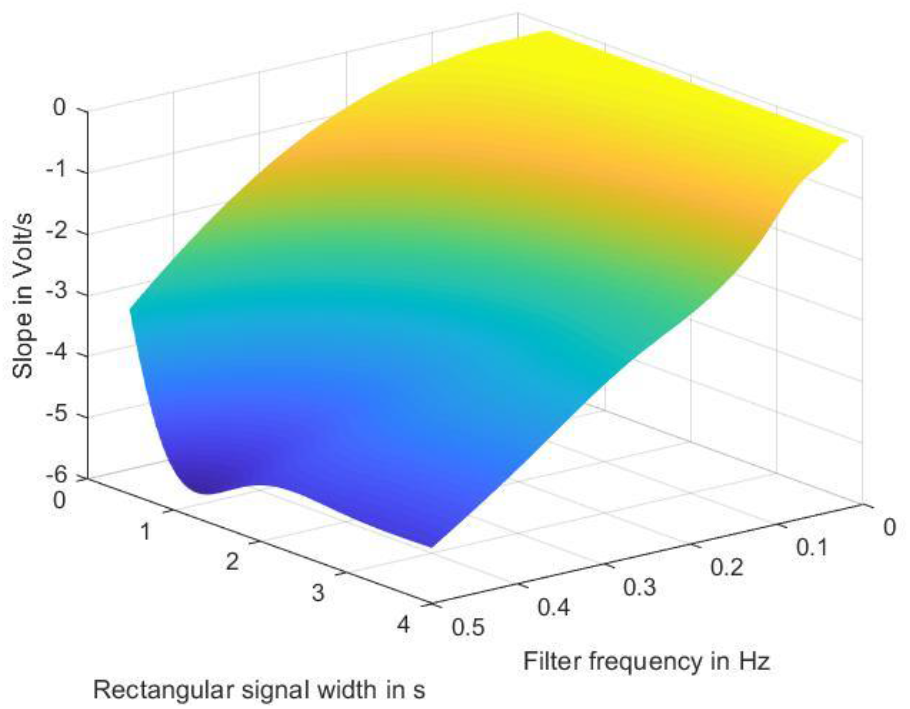
Influence of the rectangular signal width and the filter frequency on the filter error

### Part 2: Application to EEG data

To investigate the model predictions on real data, Event-Related Potentials (ERPs) and the effect of filters on them were investigated in a dataset containing task-relevant eye-movements.

## Method

### Participants and procedure

Datasets were derived from an ethically approved (UCL ethics committee ref. number: CPB/2013/011 and CPB/2014/007) study by Kulke et al. (2016a), on 23 healthy adults with normal vision (19 female, age: *M* = 21.3, *SD* = 2.4). An overt attention shift task based on the Fixation Shift Paradigm (e.g. Hood & Atkinson, 1993; Kulke et al., 2015) was used in which subjects were instructed to shift attention from a central dot subtending 0.7 degree visual angle, which was presented for a randomized inter-trial interval between 0.5 and 2.5 seconds to peripheral target bars subtending 3.1° × 13.2° randomly appearing on the left, right or on both sides of the screen at an eccentricity of 12.9°. Subjects were instructed to shift attention either by making an eye-movement towards one peripheral target (eye-movement condition containing task-relevant eye movements) or by pressing a button corresponding to one side on which the peripheral targets appeared while inhibiting any eye-movements (manual response condition). A gaze-contingent algorithm was used to control the stimulus display. Response mode (manual or saccadic) was manipulated within subjects. For the purpose of the current study, all factors and preprocessing methods were kept constant and only high-pass (HP) filter settings were varied within subjects and applied to the data set, using HP filters of 0.2, 0.1, 0.01 and 0 Hz.

### Equipment

A Tobii X120 remote eye-tracker recorded the gaze-position of subjects at a rate of 60Hz. The average viewing distance was 65 cm. Subjects completed a standard 6 point calibration routine. Participants’ manual responses were monitored with a joypad (Saitek USB V pad). EEG activity was recorded at a rate of 250 Hz using Electrical Geodesics Inc. NetAmp300 amplifier and 128-channel Ag/AgCl electrode Geodesic Sensor Nets (Tucker, 1993) on a separate computer (Macintosh) using Net Station 4.2 (© 1994-2006, Electrical Geodesics, Inc.). Electrode impedance was adjusted to less than 90 kΩ, with the majority of electrodes having an impedance of less than 40 kΩ (Ferree, Luu, Russell, & Tucker, 2001).

### Data processing

The EEG data was pre-processed with an algorithm that differed only in the high-pass (HP) filter settings. HP filters of 0.2, 0.1, 0.01 and 0 Hz were compared^1^. An average reference was used. Eye-tracking data, processed as described by Kulke et al. (2016a), was used to exclude trials with early eye-movements, ensuring that EEG data in the extracted time-windows was not confounded with eye movement artefacts. The EEG data analysis was programmed in MATLAB, using the following steps: (1.) For filtering, a 4^th^ order band reject filter was used for notch filtering around the line noise frequency [49 to 51 Hz], 3^rd^ order Butterworth filters were used for high-pass filtering using the above mentioned cut off values and low-pass filtering (fixed cut off: 25 Hz). (2.) Segmentation of data into epochs of −200 to 180 msec around target onset. (3.) Noisy epochs and electrodes were determined by using the median absolute deviation about the median (MAD) (Hampel, 1974) of amplitudes, as this is a measure that is fairly robust to noise (Hampel, 1974; Leys, Ley, Klein, Bernard, & Licata, 2013). Epochs were included if the following criteria were within this threshold calculated using the individual data in more than 70 channels: (a) SD over samples per trial, (b) range of amplitude between minimum and maximum amplitude value within epoch, (c) drift (difference in amplitude between the average period before and after target onset), (d) Maximum steps in data amplitude between successive samples; and if (e) the SD over samples per trial was bigger than 0.1 (i.e. the electrode received a signal). (4.) Other epochs were interpolated using spherical spline interpolation. (5.) The average voltage during the baseline period ([−200; 0] msec before target onset) was used to correct the data from target onset onwards, individually for each trial and electrode.

Frontocentral electrodes between electrode locations F and FC in the 10-10 system were defined as areas of interest for ERP wave plots, as these areas are both relevant for attention responses and strongly affected by eye movement artefacts (e.g. Harter, Aine, & Schroeder, 1982; Heinze, Luck, Mangun, & Hillyard, 1990; Rugg, Milner, Lines, & Phalp, 1987). Linear mixed models were computed in IBM SPSS Statistics software (version 25).

## Results

### Effects of different filter settings

Wave plots of the fronto-central responses measured in lateral electrodes in different filter conditions are displayed in Figure 3. For tests with manual responses, wave forms appear similar across different filter settings suggesting that filters do not cause significant distortion of ERP data before covert attention shifts. However, for eye movement conditions an artifactual early response in fronto-central areas is visible in the highest filter setting of 0.2 Hz, being positive contralateral and negative ipsilateral to the direction of the eye-movements. It disappears for lower HP filter settings. To statistically investigate differences between filter conditions, a linear mixed model including subject ID as random effects and response type (eye-movement or manual), and hemisphere (ipsi or contralateral) as fixed factors on mean amplitude between 80 and 120 msec was computed for each filter setting separately. Results (Table 1) show main effects of response type and hemisphere for all filter settings but the interaction is only significant for the 0.2 Hz filter.

**Table 1.** Effects for response type, hemisphere and their interaction for different filter settings.

| Filter | Response type | Hemisphere | Interaction |
| --- | --- | --- | --- |
| 0 Hz | $F(1, 3492) = 15.12,$<br>$p < .001$ | $F(1, 3492) = 27.02,$<br>$p < .001$ | $F(1, 3492) = 0.57,$<br>$p = .452$ |
| 0.01 Hz | $F(1, 3069) = 4.29,$<br>$p = .038$ | $F(1, 3069) = 19.59,$<br>$p < .001$ | $F(1, 3069) = 3.70,$<br>$p = .054$ |
| 0.1 Hz | $F(1, 2917) = 13.58,$<br>$p < .001$ | $F(1, 2917) = 9.56,$<br>$p = .002$ | $F(1, 2917) = 0.30,$<br>$p = .585$ |
| 0.2 Hz | $F(1, 2990) = 52.36,$<br>$p < .001$ | $F(1, 2990) = 91.56,$<br>$p < .001$ | $F(1, 2990) = 114.47,$<br>$p < .001$ |

**Figure 3.**
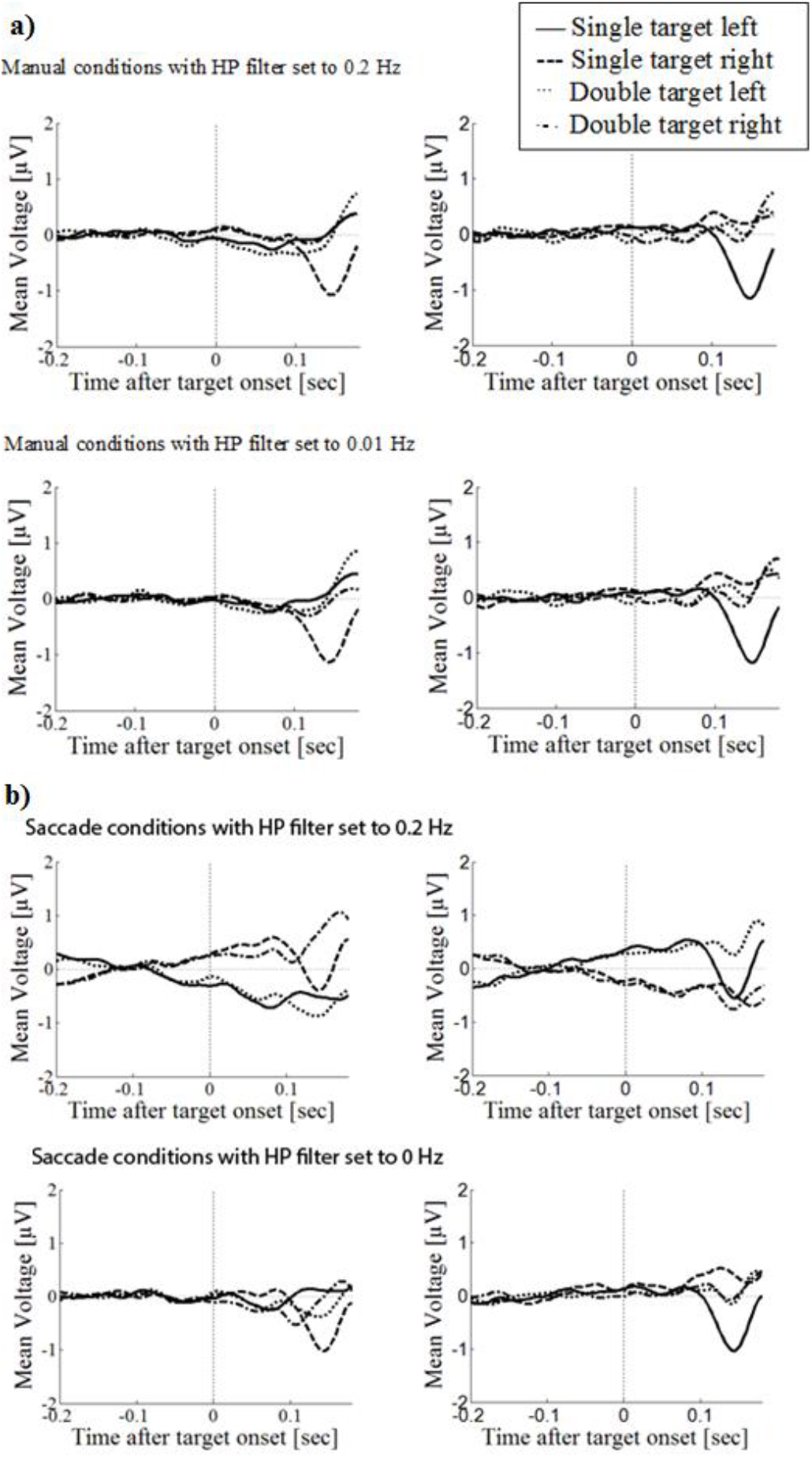
Wave plots of the fronto-central response for manual response (a) and eye-movement (b) conditions in the left hemisphere (left) and the right hemisphere (right) of the brain.

To follow up the effects in the eye-movement condition, a linear mixed model including subject ID as random effects and hemisphere (ipsi or contralateral) as fixed factors on mean amplitude was computed for each filter setting separately for the eye-movement condition only. At a *filter setting of 0* and *0.01*, mean amplitudes were significantly larger in the ipsilateral than in the contralateral hemisphere. However, at a *filter setting of 0.1*, the difference was no longer significant and at a *filter setting of 0.2*, the pattern was in the opposite direction (Table 2).

**Table 2.** Mean amplitudes in the ipsilateral and contralateral hemisphere and difference effect.

| Filter | Ipsilateral amplitude | Contralateral amplitude | Difference effect |
| --- | --- | --- | --- |
| 0 Hz | $M = -0.03$,<br>$SD = 2.28$ | $M = -0.20$,<br>$SD = 2.27$ | $F(763) = 22.95$,<br>$p < .001$ |
| 0.01 Hz | $M = -0.04$,<br>$SD = 2.458$ | $M = -0.20$,<br>$SD = 2.47$ | $F(884) = 20.86$,<br>$p < .001$ |
| 0.1 Hz | $M = -0.21$,<br>$SD = 2.17$ | $M = -0.26$,<br>$SD = 2.20$ | $F(1038) = 1.28$,<br>$p = .258$ |
| 0.2 Hz | $M = -0.57$,<br>$SD = 1.84$ | $M = 0.13$,<br>$SD = 1.93$ | $F(1713) = 233.59$,<br>$p < .001$ |

### Statistical analysis of the artefactual response

The artefactual response that was induced when using a 0.2 Hz high pass filter was further analysed by calculating peak amplitudes in different electrode locations (frontal (F3, F4), fronto-central (FC3, FC4), central (C3, C4), centro-parietal (CP3, CP4) and parietal (P3, P4)) in a time window between 80 and 120 msec that was affected by the artefactual response. Linear mixed models showed a significant effect of brain hemisphere, *F*(1, 1144) = 38.79, *p* < .001, response type, *F*(1, 1144) = 6.28, *p* = .012, a significant interaction effect of electrode location and hemisphere, *F*(4, 445) = 2.63, *p* = .034, and a significant interaction of hemisphere and response type, *F*(1, 1144) = 41.51, *p* < .001 on peak amplitude. Peak amplitudes were greater in the contralateral hemisphere (*M* = 0.83 μV, *SD* = 1.85) than in the ipsilateral hemisphere (*M* = −0.07 μV, *SD* = 1.53). Figure 4 shows that the hemispherical difference is greater in frontal areas and decreases towards parietal areas.

**Figure 4.**
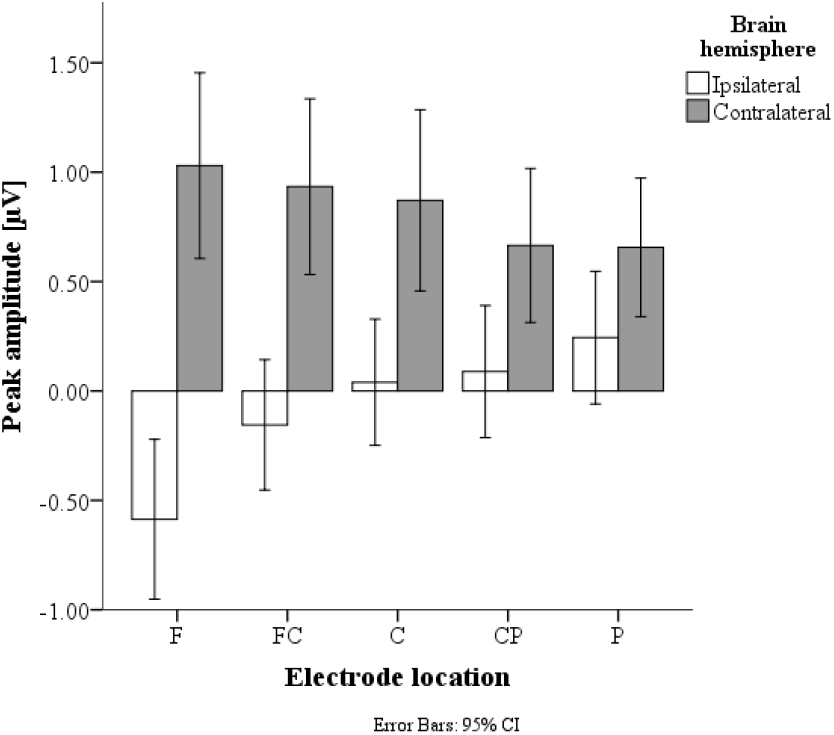
Artifactual responses are more lateralised in frontal regions (F) and decrease towards parietal regions (P).

### Time window before stimulus onset

To further explore the onset of the fronto-central response at the 0.2 Hz filter setting, the mean amplitude in the time window before stimulus onset from −50 to 0 ms in the eye-movement condition was analysed with a linear mixed model with participants as random effects and number of targets, brain hemisphere and brain side as fixed effects. There was a significant effect of hemisphere on mean amplitude, *F*(1, 85) = 5.40, *p* = .023.

## Discussion

The analyses show that filters can induce artefactual responses in EEG data containing regular eye-movements. A clear artefactual response was identified if a 0.2 Hz HP filter was used, resulting in an opposite polarity compared to effects observed with lower filter settings, and a decreased ipsilateral and increased contralateral amplitude, i.e. the opposite polarity to an eye-movement, as expected as a result of excessive filtering (Tanner et al., 2015). Although no significant interactions of response type and hemisphere were observed for 0.1 Hz filters, the original effect observed using lower filter settings (Kulke et al., 2016a) disappeared at 0.1 Hz, suggesting that this criterion already affects findings.

Follow-up analyses of the artefactual response showed the artefacts had a statistically significant effect on ERP measurements. Detailed analyses support the artificial nature of the response, showing that the measured slow wave is more lateralised in anterior areas, which are more affected by eye-movements, than in posterior areas. It was furthermore already significantly lateralised before target onset, suggesting that it is unrelated to visual input, as, by definition, no visually evoked potentials can occur before the visual event.

Given that high-pass filtering can induce artefactual responses due to eye-movements, the high-pass filter criterion should be set to a minimum to avoid artefacts. Tanner et al. (2015) suggested that for EEG data without any task-relevant eye-movements, high pass filters below 0.3 Hz should be used. However, the current study shows that with this criterion artefacts are already induced in data containing task-relevant eye-movements.

A recent publication by Dimigen (2020) investigates improvements of EEG data quality through ICA for co-registered EEG and eye-tracking data sets. It demonstrates that the removal of artefacts through ICA is improved when stronger HP filters are used; however, with stronger filters the overcorrection at frontal sites also increases, leading to larger distortions of the data. This finding is in line with our models, suggesting distortions due to strong HP filters. Filter settings may therefore even play a role if additional artefact removal is conducted using ICA.

In conclusion, studies containing task-relevant eye movements are prone to filter artefacts induced by high-pass filtering and cut-off criteria should therefore be set lower than 0.1 Hz unless further corrections to the data distortions can be made.

## Acknowledgments

The research was funded by a German Academic Exchange Service stipend to LK [DAAD, grant number 91522396-57044644]. The EEG analysis was based on programs written by John Wattam-Bell. We would like to thank Ankita Agharkar and Megan Gawryszewski for their help with data collection, Jyrki Toumainen for helpful feedback on EEG data analysis and Janette Atkinson and Oliver Braddick for helpful feedback regarding the manuscript.

## Supplementary figures

**Figure S.1.**
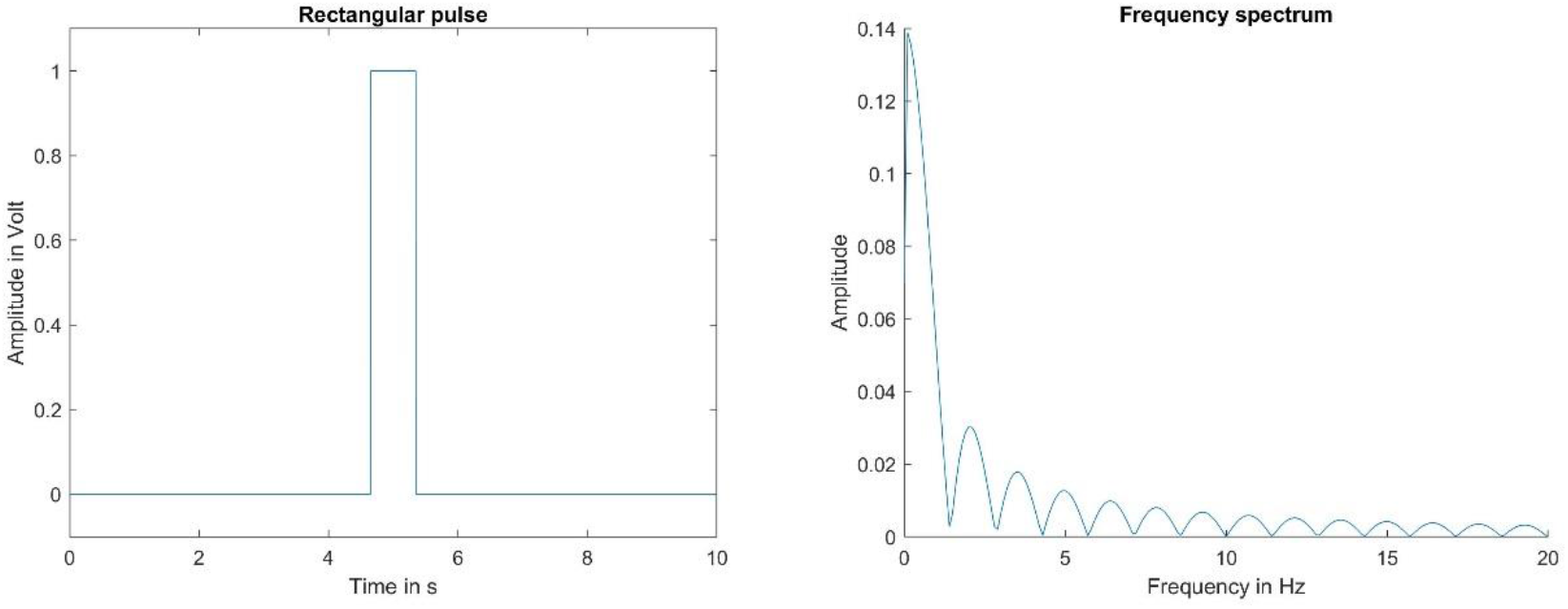
Rectangular signal and its associated frequency spectrum

**Figure S.2.**
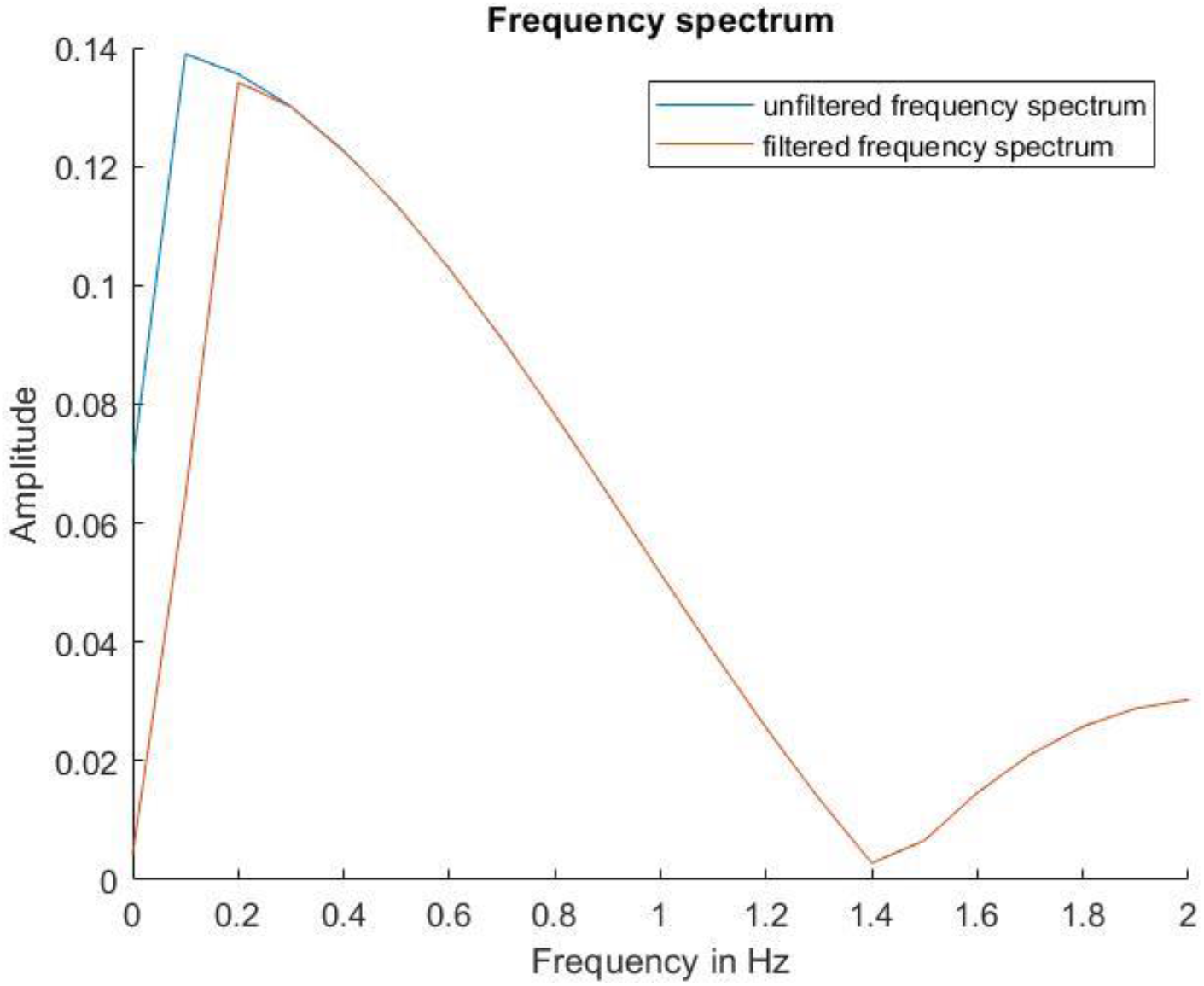
Frequency spectrum of a filtered and an unfiltered rectangular pulse

**Figure S.3.**
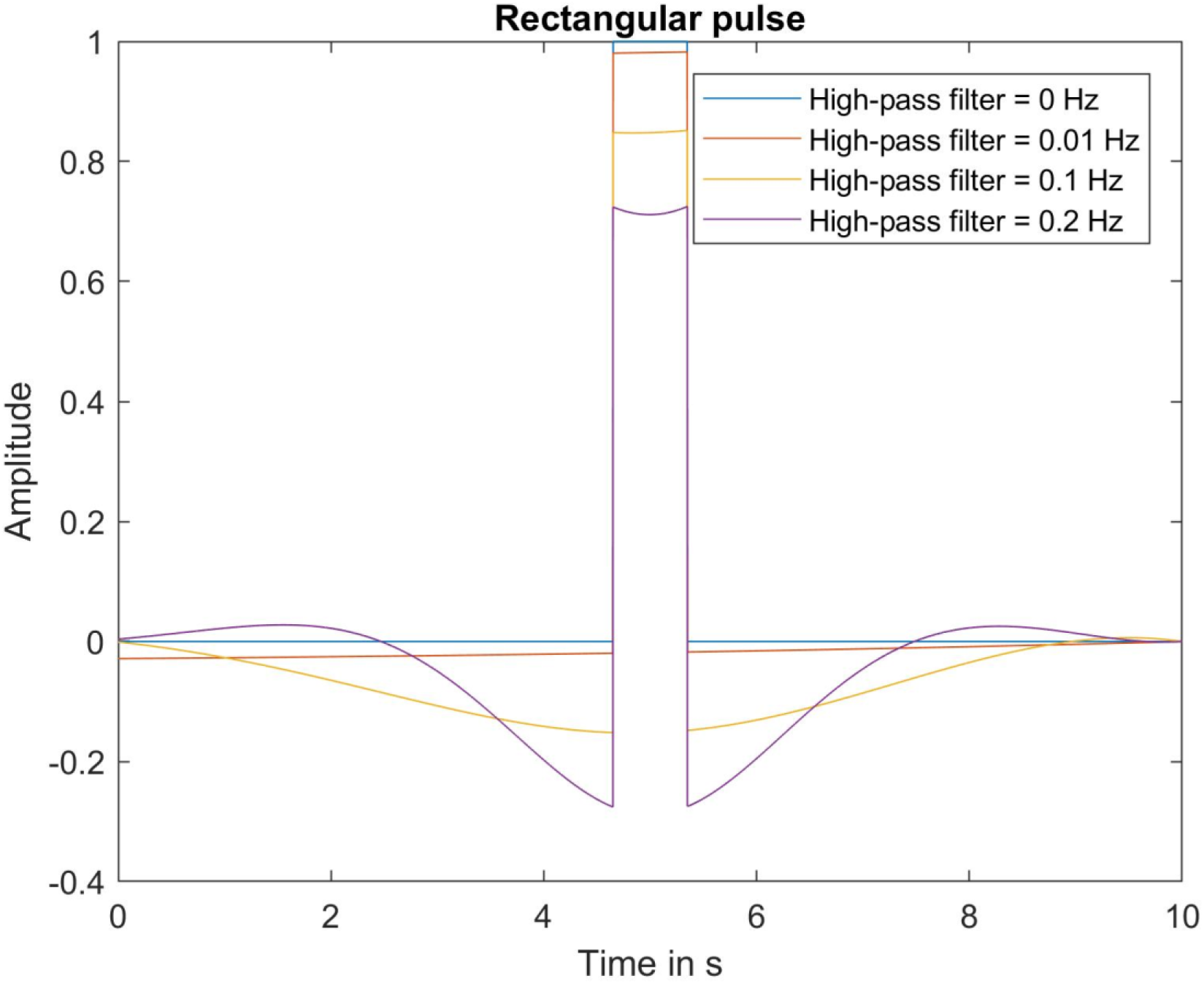
Time signal of a rectangular pulse with the filter error after inverse FFT

**Figure S.4.**
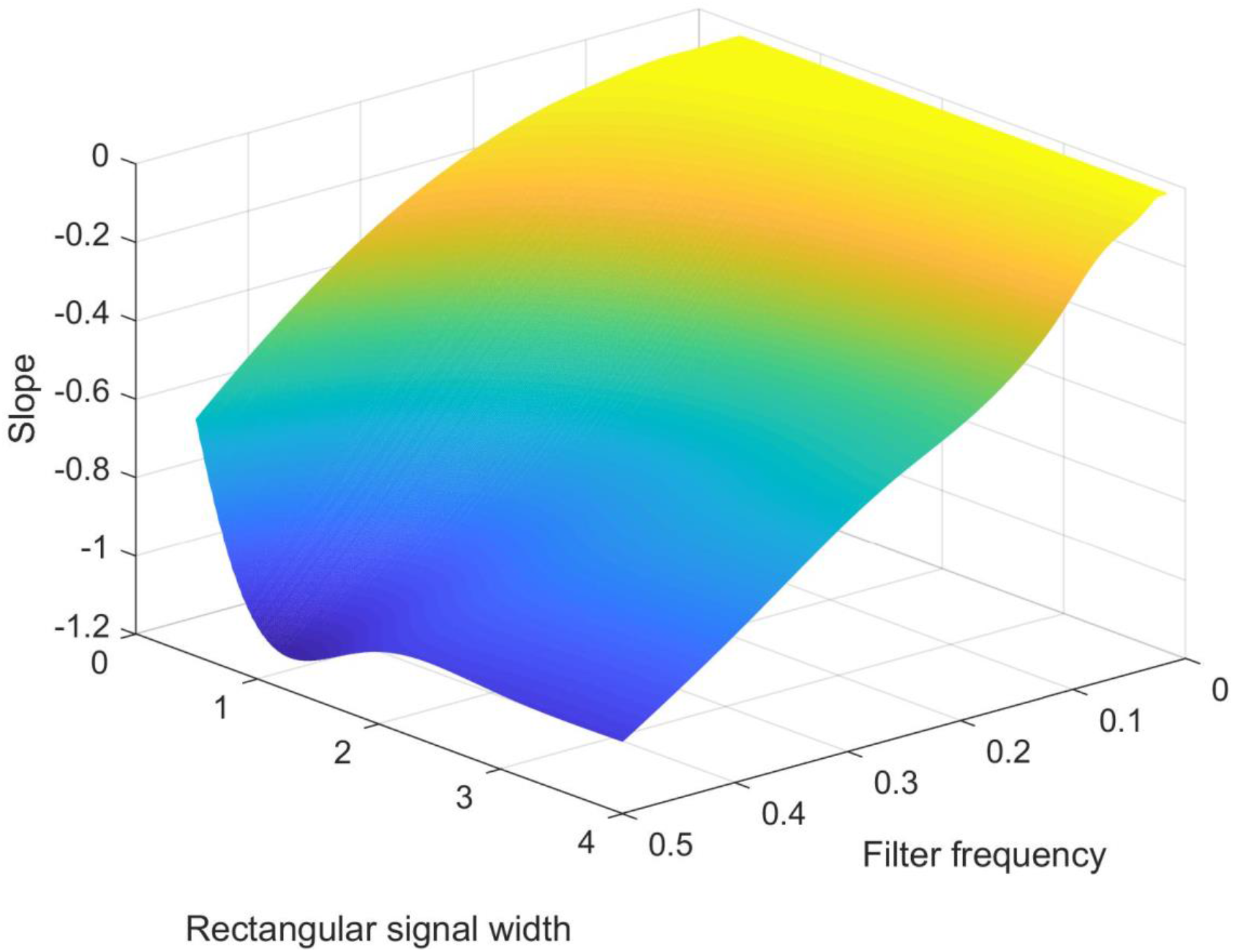
Influence of the rectangular signal width and the filter frequency on the filter error

**Figure S.5.**
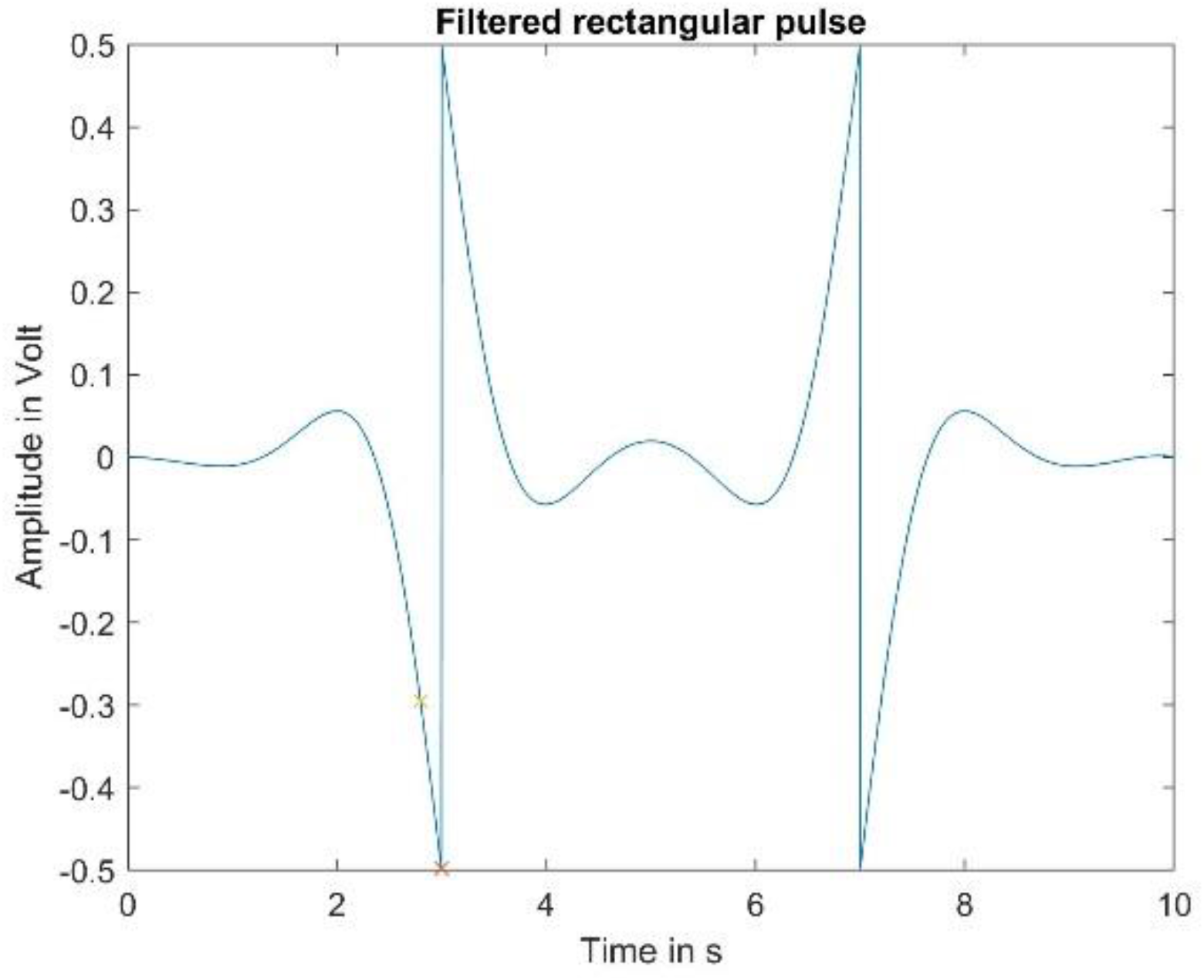
Highly distorted rectangular pulse due to strong filters and long pulse width

## Footnotes

1 The NetAmp 300 amplifier automatically applies an analogue low-pass filter at 6 kHz and after the analogue-digital conversion of the data applies a 4 kHz low-pass filter. No further online high-pass or low-pass filters were applied to avoid data distortion.

